# Design and validation of a PCR protocol to specifically detect the clade of *Philaster* sp. associated with *Diadema antillarum* scuticociliatosis

**DOI:** 10.1101/2023.09.11.557215

**Authors:** Brayan Y. Vilanova-Cuevas, Brandon Reyes-Chavez, Mya Breitbart, Ian Hewson

**Affiliations:** Department of Microbiology, Cornell University, Ithaca NY USA; College of Marine Science, University of South Florida, St Petersburg FL USA

## Abstract

*Diadema antillarum* scuticociliatosis (DaSc), caused by a scuticociliate closely related to *Philaster apodigitiformis*, has affected Caribbean long-spined urchins since at least January 2022. Quantitative PCR (qPCR) is currently the standard method for detection of this ciliate in tissue and coelomic fluid samples, yet this method requires specialized equipment and is more expensive than standard PCR methods. The DaSc scuticociliate occurs against a backdrop of endo- and ecto-symbiotic ciliates which complicate detection using universal or pan-phylum PCR primer sets. To overcome these limitations, we designed and validated a sensitive and specific PCR primer (scutico-634F) and nested two-step PCR protocol to detect this taxon, which excludes other ciliates associated with *D. antillarum* and has poor affinity for other related ciliates. This primer and protocol for the DaSc-associated Philaster clade (DaScPc) allow for widely-accessible investigation of this pathogen in new regions and within environmental reservoirs.

## INTRODUCTION

Ciliates (Alveolata: Ciliophora) are ecologically and biogeochemically significant protistan inhabitants of aquatic habitats worldwide (Hausmann & Bradbury 1996). They are important predators of bacteria, archaea and other protists, and thus play key roles in aquatic food webs. Ciliates are also known to form symbioses with a variety of aquatic metazoans, with interactions ranging from commensalism to parasitism (Rosaura et al. 2021). Ciliates inhabiting sea urchins have been studied for over a century (Hoffmann, 1871; Jacobs 1914) and are known to form ectocommensal and endosymbiotic relations that play necessary roles in their gut where they may break down complex food (Beers 1961, Jangoux 1987, Wells 2013; Table 1). While many ciliates are described as commensals of echinoderms, they can also cause disease or serve as opportunistic pathogens in response to environmental stressors (Bouland & Jangoux 1988, Cawthorn et al. 1996).

**Table 1.** Summary of reported commensal ciliates that inhabit *Diadema spp*. urchins.

| *Diadema*<br>species | Ciliate taxon | Geographic<br>Location | Reference |
| --- | --- | --- | --- |
| *D.*<br>*antillarum* | *Cohnilembus caeci*,<br>*Biggaria bermudensis*, *Cyclidium*<br>*rhabdotectum*, *Metopus*<br>*circumlabens*, *Metopus rotundus*,<br>*Anophrys aglycus*, *Anophrys elongata*,<br>*Dysteria* sp., *Cryptochilidium*<br>*bermudense* | Caribbean,<br>Bermuda | Jacobs 1914, Biggar &<br>Wenrich 1934, Lucas<br>1934, Powers 1935,<br>Uradaneta-Morales &<br>de McLure 1966, Jones<br>& Rogers 1968 |
| *D.*<br>*savignyi* | *Cryptochilidium polynucleatum*,<br>*Metopus circumlabens* | American<br>Samoa | Anderson 1972 |

The long spined Caribbean urchin, *Diadema antillarum* experienced mass mortality across the Caribbean starting in spring 2022 (Hylkema et al. 2023). This mass mortality echoed an earlier event which decimated *D. antillarum* population densities from 1983-1984 and triggered a phase shift from coral to algal-dominant environments in the region (Lessios 2016). While the cause of the 1980s mass mortality was never identified, the 2022 event was caused by a scuticociliate most closely related to *Philaster apodigitiformis* based upon 28S rRNA sequence (Hewson et al. 2023) and the disease has been named *Diadema antillarum* scuticociliatosis (DaSc). Scuticociliatosis, caused by ciliates from the Scuticociliatia subclass, is one of the most diverse and widely studied aquatic diseases affecting a diversity of hosts (Azad et al. 2007). Commensal ciliates are common within the gut of *Diadema* spp. (Table 1). Unlike other previously described commensal organisms that are restricted to the digestive tract of *Diadema antillarum*, this scuticociliate taxon is found in several tissue types and coelomic fluid (Hewson et al. 2023).

The DaSc-associated *Philaster* occurs in affected hosts amongst a backdrop of other ciliate taxa, which complicates detection through molecular methods. Universal ciliate or pan-phylum PCR primer sets therefore cannot be used to specifically detect the pathogen of interest in sea urchins. Furthermore, the quantitative PCR (qPCR) approach employed by Hewson et al. (2023) may not be accessible to labs without qPCR instruments, and the cost of reagents and fluorophore tagged primers may prohibit broad application of this approach. Since the 2022 Caribbean mortality event, additional mass mortalities have been described in other geographic regions and in related *Diadema* species (Zirler et al. 2023). To enable simple and specific detection of the DaSc-associated *Philaster* clade for diagnostic purposes, here we report a new PCR primer and amplification protocol that selectively amplifies the DaSc-associated *Philaster*. We validated this primer by amplifying specimens known to contain the scuticociliate using general ciliate primers and assessed specificity by comparison against sequences of other ciliates recovered from *Diadema antillarum*.

## METHODS

### Echinoderm tissue sampling

Spine, body wall, and coelomic fluid samples were obtained as part of a prior effort to understand the etiology of the 2022 Caribbean *Diadema antillarum* scuticociliatosis outbreak (Hewson et al. 2023). DNA was extracted using the Zymo Tissue and Insect DNA kit following manufacturer’s recommendations as reported in Hewson et al. (2023). For each sampling location, equal extraction volumes of DNA were by tissue type (coelomic fluid, body wall, and spine+spine base) for each health state (reference, normal at affected site, and DaSc-affected), resulting in 53 sample pools (Table 2).

**Table 2:**
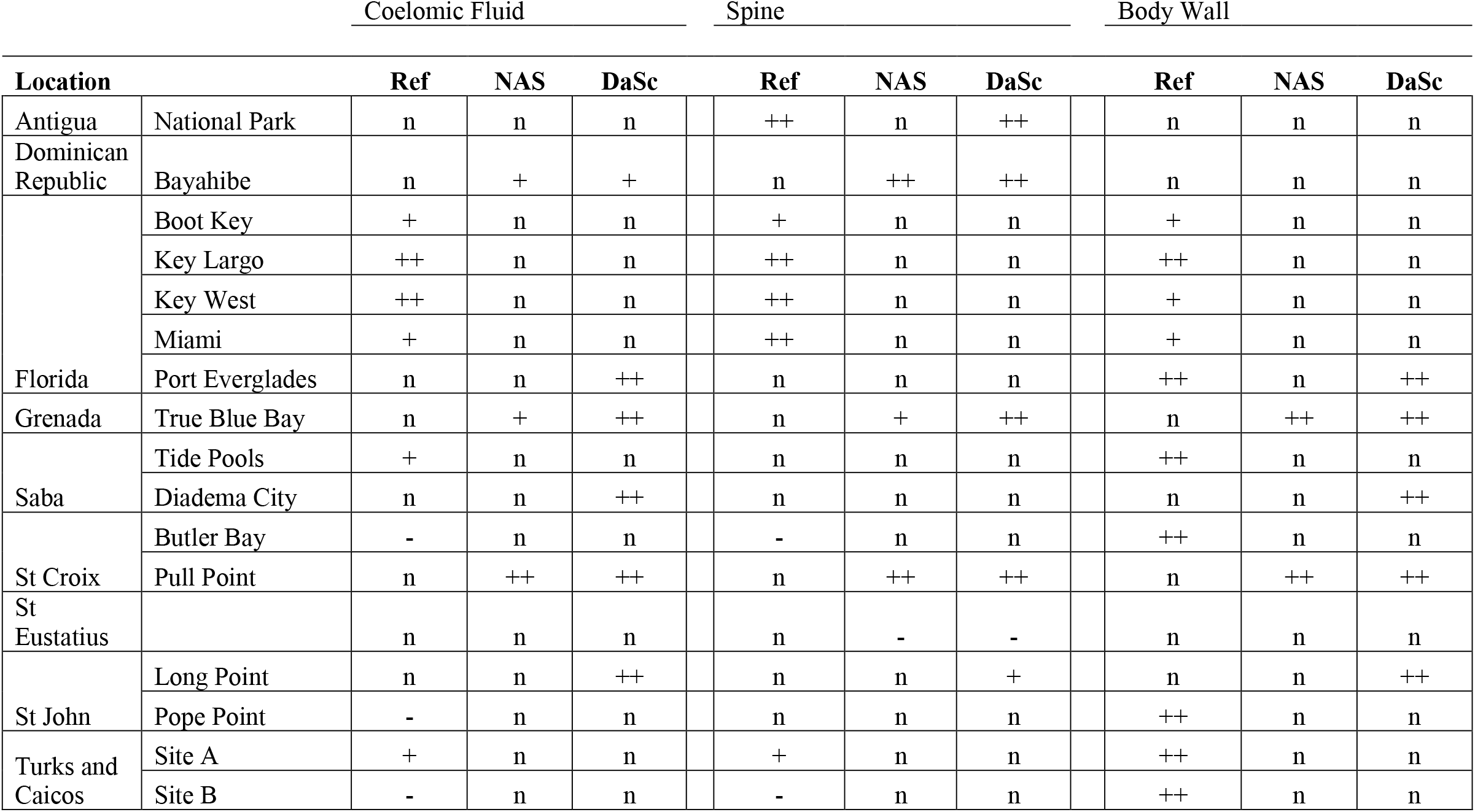
Sampling locations and intensity of amplicon on electrophoretic gels using the pan-Ciliophora primers 387F and 1147R (Dopheide et al. 2008). Note that not all positive amplicons generated PCR sequences via Sanger sequencing, which presumably was due to insufficient DNA concentrations for sequencing chemistry. + = faint amplicon, ++ bright amplicon, - no amplicon, n = sample not examined, Ref = Reference Site, NAS = grossly normal urchin at DaSc-affected site, DaSc = DaSc-affected urchin.

### PCR amplification with general primers

Pooled DNA extracts were subjected to PCR amplification using primers 384F (5’ - YTB GAT GGT AGT GTA TTG GA - 3’) and 1147R (5’-GAC GGT ATC TRA TCG TCT TT - 3’) as described by Dopheide et al. (2008) targeting the 18S rRNA gene of Ciliophora. PCR was performed in 50 μl reactions containing 1X PCR Buffer (ThermoFisher), 2.5 mM MgCl_2_, 0.2 mM PCR Nucleotide Mix (Promega) 10 μM of each primer, 5U Taq DNA polymerase (ThermoFisher), 0.02 ng μl^-1^ BSA, and 2 μl of template DNA. PCR reactions were subjected to an initial heating step at 95°C for 3 min, followed by 30 cycles of denaturation at 95°C for 30 s, annealing at 54°C for 30 s, and extension at 72°C for 30 s. Thermal cycling was concluded with a final extension step at 72°C for 5 min.

PCR products and a standard ladder (Promega) were electrophoresed on 1% agarose gels in 1X TBE for 1 hr at 85V. After staining gels with SYBR Gold (10X), gels were visualized on a BioRad ChemiDoc system. Reactions with visible products at ∼750 bp were purified using the Zymo Clean & Concentrator −5 kit and submitted to the Cornell University Biotechnology Research Center for Sanger sequencing using the 384F primer. All sequences included in this study are available in GenBank under accession numbers OR525708-OR525738.

Sequences were first trimmed to remove ambiguous base calls at the 5’ and 3’ sequence ends, then subjected to BLASTn (Altschul et al. 1997) against the NCBI nonredundant (nt) database current as of December 2022. The best match for each sequence based on lowest e-value was then assembled into a single fasta file, which was aligned with the products from this study using the MUSCLE algorithm (Edgar 2004). After masking to a contiguous region, evolutionary history was inferred using the maximum likelihood method in MEGAX (Kumar et al. 2018). The optimal model of nucleotide substitution was determined by the highest Bayesian information criterion (BIC) value, which identified the Tamura 3 parameter with gamma distributed sites. A phylogenetic tree was generated based on 100 bootstrap iterations.

### Design of DaSc-associated Philaster primer

The phylogenetic analysis performed on *Diadema*-associated ciliates enhanced phylogenetic resolution compared to previous analyses (Hewson et al. 2023) and revealed a clade of 18S rRNA sequences that were primarily found in DaSc-affected urchins. This clade includes cultures FWC1 and FWC2 (both cultivated from field-collected urchins in Key Largo, Florida), USF152 (re-isolated from urchins challenged with FWC2 that developed DaSc), as well as an 18S rRNA gene sequence assembled from DaSc-affected specimens from Saba and St John. We term this clade DaSc-*Philaster* clade (DaScPc).

Because this clade is distinct from the 18S rRNA sequences of *Philaster lucinda* and *Philaster guamensis* (both recovered from corals; Sweet & Bythell 2012, Sweet & Séré 2016, Sweet 2020) and *Philaster apodigitiformis* (originally identified in from pufferfish eggs; Miao et al. 2009), future reference to DaSc-associated ciliates should follow this clade designation until formal taxonomy is applied. We then sought to design a primer that specifically amplifies DaScPc to enable rapid identification of the infectious agent for DaSc. The DaScPc comprises urchin-derived sequences that were 99-100% identical to the FWC2, FWC1 and USF152 cultures, so we used their 18S rRNA sequences in downstream design.

Using MUSCLE, we created an alignment with the DaScPc sequences, the best BLASTn matches obtained from urchins using the general ciliate primers, and representative 18S rRNA gene sequences spanning the Ciliophora. We manually examined conserved nucleotide sequence regions and found a region within the amplicon obtained with the general ciliate primers (384F/1147R; (Dopheide et al. 2008)) on which to base the DaScPc primer. We then used the Primer3 program (Rozen & Skaletsky 2000) to identify a primer sequence (scutico-634F; 5’ - TTG CAA TGA GAA CAA CGT AA - 3’) that selectively amplified DaSCPc sequences over related taxa. BLASTn analysis of the primer revealed that it shared 100% sequence identity with some metazoans but was not similar to other ciliates and protists in GenBank. We validated this new scutico-634F primer in combination with 1147R from Dopheide et al. (2018) with the same PCR conditions described above except lowering the annealing temperature to 53°C, using 1 μl of the product from the general ciliate PCR as template (i.e., nested PCR). PCR was performed on three specimen pools which generated DaScPc amplicons with the general ciliate primers.

## RESULTS AND DISCUSSION

A total of 53 *Diadema antillarum* specimen pools yielded amplicons with the general ciliate primers, however only 35 had sufficient DNA quantities for Sanger sequencing of the PCR products. Of these, 28 yielded sequences (the rest failed due to insufficient amplicon quantity), suggesting that this approach may only be successful when ciliate densities are high enough to yield sufficient template. Of these 28 sequences, 15 sequences fell within DaScPc, with 12 sequences originating from abnormal specimens and 3 sequences from grossly normal specimens at affected sites (Fig. 1). All three tissue types (8 coelomic fluid, 8 body wall and 12 spine pools) yielded DaScPc sequences, which is consistent with quantitative PCR results based on the 28S rRNA locus (Hewson et al., 2023). DaScPc sequences were most often recovered from Grenada (4 sequences from 6 pools), Florida (8 sequences from 15 pools) and St Croix (5 sequences from 9 pools) and least in the Turks and Caicos Islands (1 sequence from 6 pools). The next most common ciliate sequence detected was most similar to *Licnophora macfarlandi* (1 abnormal spine pool, 4 grossly normal pools at affected sites and 1 at a reference site), followed by sequences most similar to *Parametopidium circumlabens* (1 normal spine pool at an affected site and 3 spine pools from reference sites). A single reference sequence matched *Biggaria bermudensis*.

**Fig. 1:**
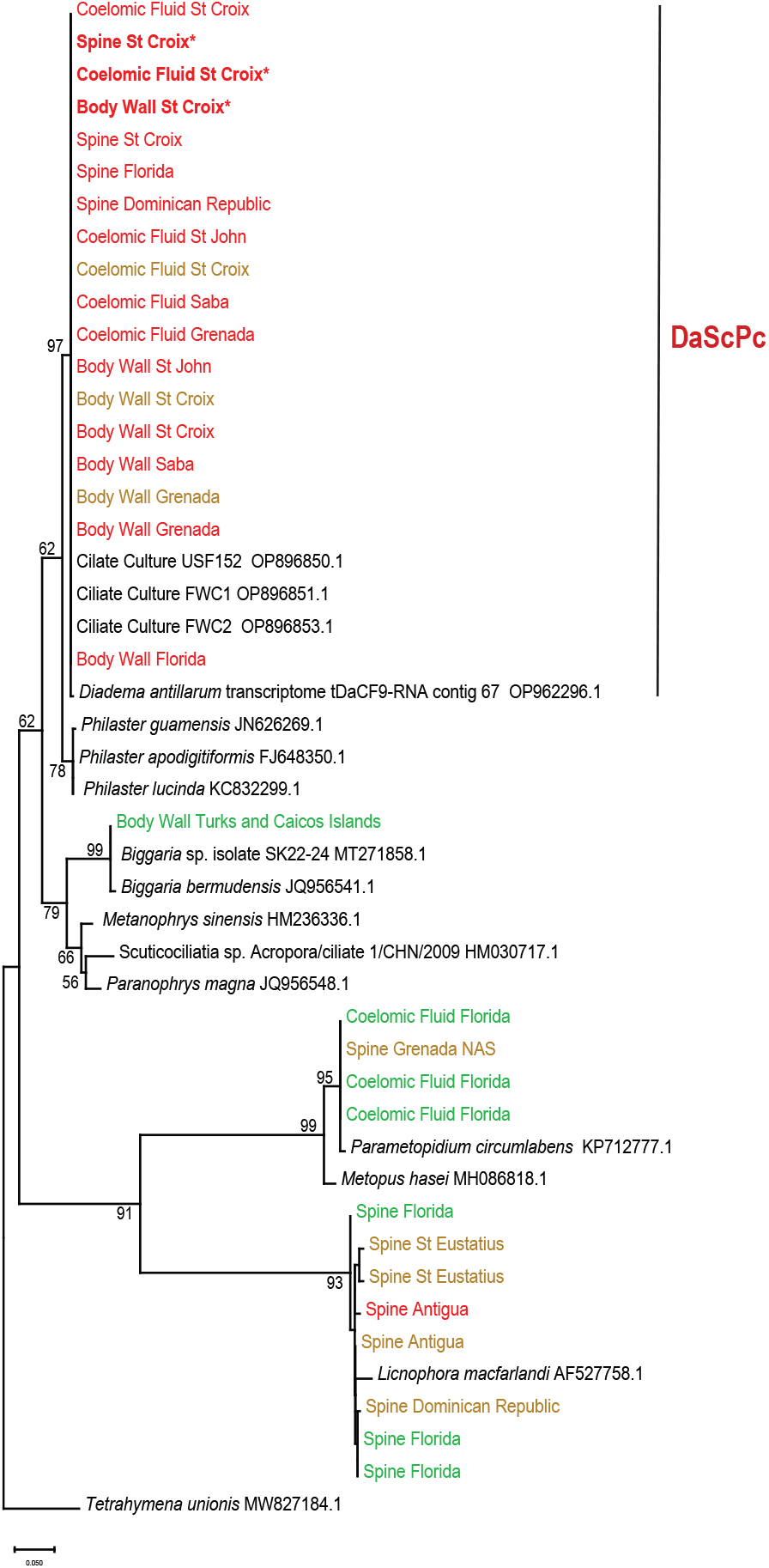
Phylogenetic reconstruction of *Diadema antillarum*-derived 18S rRNA sequences and close relatives as identified by BLASTn searches of the nonredundant (nr) database. The evolutionary history was constructed in MEGAX (Kumar et al. 2018) and inferred by Maximum Likelihood and the Tamura-3 parameter model and is based on neighbor joining. Bootstrap values represent confidence in 100 iterations. The scale bar indicates substitutions per site. Labels marked with * indicate sequences amplified using the scutico-634F primer. The sequences are color coded to correspond with abnormal (red), grossly normal at affected site (brown) and grossly normal (green) conditions.

These results confirm previous efforts using transcriptomics and qPCR to understand the etiology of *Diadema antillarum* mass mortality (Hewson et al. 2023). The philasterine 28S rRNA sequence obtained in transcriptomes of affected urchins shared >99% nucleotide identity to the 28S rRNA of *Philaster apodigitiformis*. Strikingly, while the only ciliate 28S rRNA sequence recovered in the transcriptomes of affected urchins was nearly identical to this taxon, the 18S rRNA sequence in the same survey was only 96% identical across the locus to any *Philaster* species. These results, combined with the general ciliate 18S rRNA survey presented here, suggests that nucleotide identity between *Philaster* species may be more conserved in the 28S rRNA locus than the 18S rRNA locus. Because abnormal urchins yielded a clade of sequences that was separated with >50% bootstrap values from the *Philaster lucinda, Philaster guamensis* and *Philaster apodigitiformis* cluster, we recommend that subsequent research refers to the DaSc-associated 18S rRNA sequences as the *Diadema antillarum* Scuticociliatosis Philaster clade (DaScPc). Because of the tight association between occurrence of this sequence and gross signs of DaSc, we suggest this sequence is a sensitive biomarker of the condition.

The remaining taxa identified in this survey reflect previous observations of ciliates associated with *Diadema antillarum. Licnophora macfarlandi* has been observed in association with coral disease (Sweet & Séré 2016, Ravindran et al. 2022), and may constitute a cosmopolitan taxon in coral reef settings. *Parametopidium* spp. is a well-known, anaerobic, endocommensal ciliate in sea urchins (da Silva-Neto et al. 2016). *Biggaria bermudensis* has long been known to associated with various Caribbean sea urchin species (Biggar & Wenrich 1932). Surveys based on general ciliate 18S rRNA gene primers may identify these taxa across a wide array of urchin specimens and may dominate amplicon sequence libraries or signals when DaSc is rare during early infection.

This study demonstrated that general ciliate 18S rRNA primers may yield DaScPc sequences in abnormal urchins where DaScPc is presumably the dominant Ciliophora taxon. However, since DaScPc ciliates may be present at low abundance early in infection, and to enable DaScPc surveys of environmental reservoirs or detection via environmental DNA (eDNA) analysis where DaScPc is rare compared to other ciliate taxa, this internal PCR primer to selectively amplify DaScPc ciliates is a valuable tool for the research community. The primer has a single purine:purine mismatch near the 3’ end to *Philaster* spp., and up to 5 mismatches (primarily purine:purine mismatches) with more distantly related Scuticociliatia (including *Biggaria* which was detected in a reference urchin specimen; Table 3). Additionally, other more distantly related Ciliophora had > 3 mismatches to the primer, including at the 5’ end which may have a more significant effect under typical PCR conditions (Kwok et al. 1994). Hence, the scutico-634F primer should provide specific amplification of DaScPc over more distantly related scuticociliates or other members of the Ciliophora. The calculated primer melting temperature is 54.2°C for 634F and 57.3°C for the 1147R primer. Therefore, the annealing temperature of 53°C used in this PCR protocol should provide stringency during thermal cycling by selecting against the purine:purine mismatch found in closely related philasterine ciliates. However, if resources are available, we still recommend sequencing positive PCR products to confirm their identity, especially when applying this assay to new species or in new geographic locations.

**Table 3:**
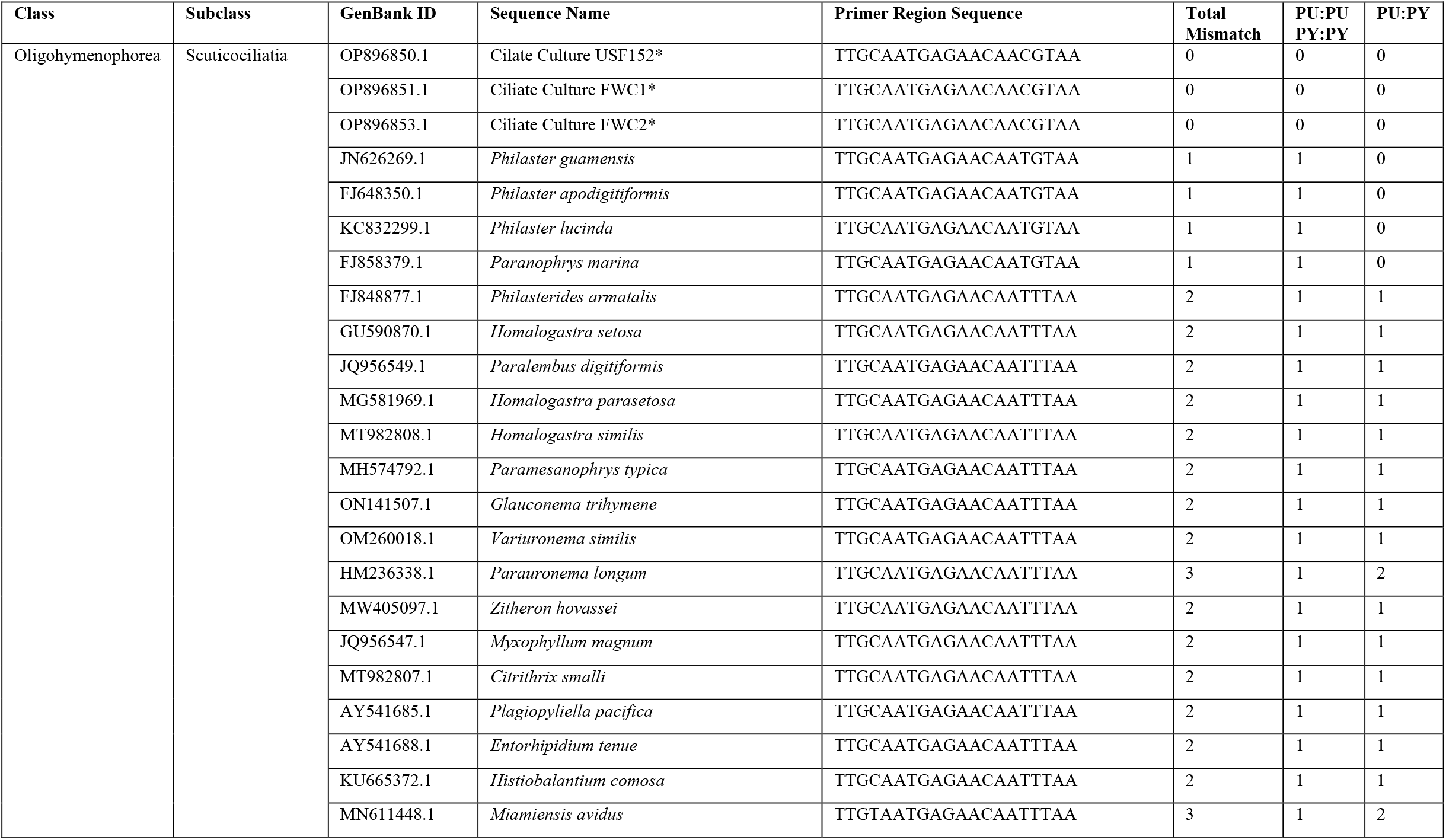
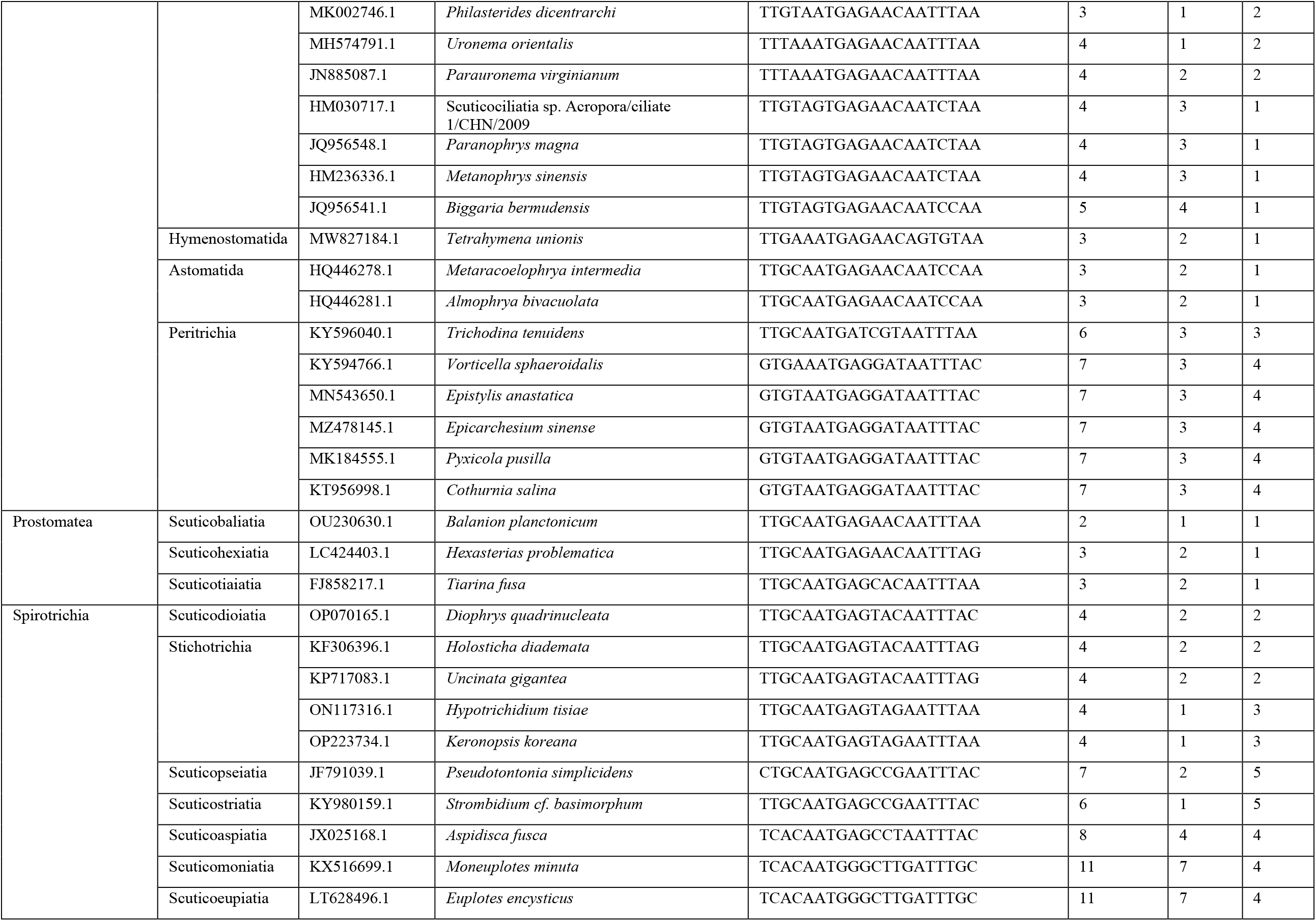
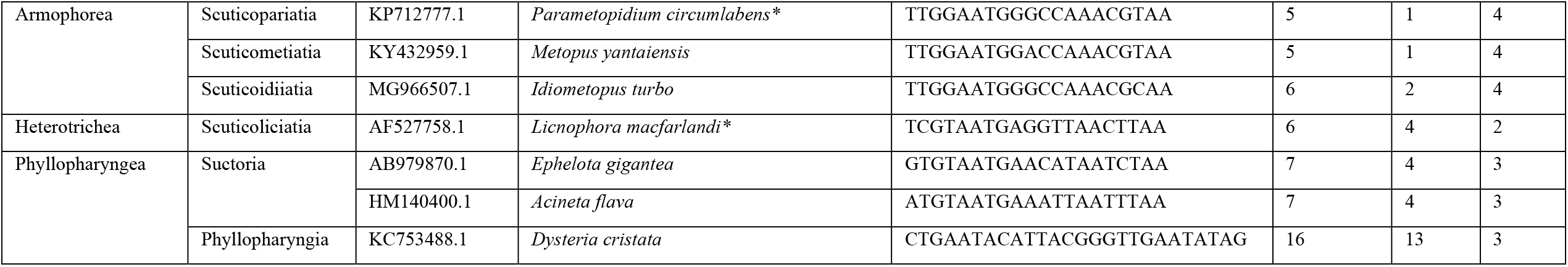
Mismatch characteristics between the priming region (matching the cultures FWC1, FWC2 and USF152). The number of purine:purine + pyrimidine:pyrimidine (G:A, C:T) and purine:pyrimidine (G:C, G:T, A:T, A:C) are also shown. Mismatch tolerance is greatest for purine:pyrimidine mismatches in primer design under stringent PCR conditions (Kwok et al. 1994). Taxa with relatives that were detected using the pan-Ciliophora primer set 384F and 1147R (Dopheide et al. 2008) in this survey are marked with an *. PY = pyrimidine; PU = purine.

The new primer and PCR protocol described in this study provides a more accessible approach for general molecular biology laboratories without specialized equipment or large budgets for qPCR. These labs may use any appropriate reagent suppliers, electrophoretic conditions for detecting amplicons, or sequencing approaches amenable to direct PCR product sequencing. Furthermore, analysis of sequences is straightforward using a publicly accessible resource (NCBI GenBank). Combined with deployable “bento box” style molecular biology platforms, this approach may also enable rapid, field-based detection of the DaSc pathogen.

## ACKNOWLEDGEMENTS

The authors are grateful to M. Pistor, M. Warham, A. Zimmermann, M. Brandt, M. Sevier, D. Marancik, A. Croquer, R. Camacho, G. Delgado and W. Sharp for collection of urchin samples from which DNA was extracted for use in this survey. This work was supported by an award from the Atkinson Center for Sustainable Futures, and NSF OCE-2049225 awarded to IH. BYVC was supported by an NSF Graduate Research Fellowship while performing this work.

